# Live imaging of *Aiptasia* larvae, a model system for studying coral bleaching, using a simple microfluidic device

**DOI:** 10.1101/370478

**Authors:** Will Van Treuren, Kara K. Brower, Louai Labanieh, Daniel Hunt, Sarah Lensch, Bianca Cruz, Heather N. Cartwright, Cawa Tran, Polly M. Fordyce

**Affiliations:** Department of Microbiology and Immunology, Stanford University, Stanford, CA 94305; Department of Bioengineering, Stanford University, Stanford, CA 94305; Department of Chemical Engineering, Stanford University, Stanford, CA 94305; Department of Genetics, Stanford University, Stanford, CA 94305; Department of Physics, California State Polytechnic University, Pomona, CA 91768; Department of Plant Biology, Carnegie Institution for Science, Stanford, CA 94305; Chem-H Institute, Stanford University, Stanford, CA 94305; Department of Biological Sciences, California State University, Chico, Chico, CA 95929; Chan Zuckerburg BioHub, San Francisco, CA 94158

## Abstract

Coral reefs, and their associated diverse ecosystems, are of enormous ecological importance. In recent years, coral health has been severely impacted by environmental stressors brought on by human activity and climate change, threatening the extinction of several major reef ecosystems. Reef damage is mediated by a process called ‘coral bleaching’ where corals, sea anemones, and other cnidarians lose their photosynthetic algal symbionts (genus *Symbiodinium*) upon stress induction, resulting in drastically decreased host energy harvest and, ultimately, coral death. The mechanism by which this critical cnidarian-algal symbiosis is lost remains poorly understood. Here, we report ‘Traptasia’, a simple microfluidic device with multiple traps designed to isolate and image individual live larvae of *Aiptasia*, a sea anemone model organism, and their algal symbionts over extended time courses. *Aiptasia* larvae are **~**100 μm in length, deformable, and highly motile, posing particular challenges for long-term imaging. Using a trap design optimized via fluid flow simulations and polymer bead loading tests, we trapped *Aiptasia* larvae containing algal symbionts and demonstrated stable imaging for >10 hours. We visualized algal migration within *Aiptasia* larvae and observed algal expulsion under an environmental stressor. To our knowledge, this device is the first to enable live imaging of cnidarian larvae and their algal symbionts and, in further implementation, could provide important insights into the cellular mechanisms of coral bleaching under different environmental stressors. The device is simple to use, requires minimal external equipment and no specialized training to operate, and can easily be adapted to study a variety of large, motile organisms.

---

Coral reefs are remarkably productive ecosystems, supporting approximately 9% of the ocean fish biomass and 25% of oceanic species diversity1. Reef-building corals depend on an endosymbiotic relationship with dinoflagellate algae (genus *Symbiodinium*) for survival and productive growth(2, 3), as most corals and other symbiotic cnidarians derive their primary metabolic energy through algal photosynthesis4. Under environmental and anthropogenic stressors such as rising ocean acidity, pollution, and increasing temperature, the coral-algal symbiotic relationship can break down, resulting in a process known as ‘coral bleaching’(1, 5). In this process, photosynthetic algal symbionts are expelled from the coral gastrodermal tissue where they normally reside, resulting in coral discoloration and, if prolonged, host death due to insufficient energy metabolism5. On a macro-scale, coral bleaching has been implicated as a major cause of rapid worldwide reef deterioration and ecosystem disruption1. To address this threat, coral-algal symbiosis, in general, and the mechanism of coral bleaching, in particular, must be better understood.

While candidate stressors that contribute to coral bleaching are known, the molecular and cellular mechanisms of algal expulsion remain poorly characterized, in part due to the difficulties of working with corals in the laboratory(3, 6). In this work, we use a small sea anemone, *Aiptasia*, as a model system for coral symbiosis(6–9). *Aiptasia* is an attractive organism for mechanistic biology studies of coral because both larval and adult animals are compatible with laboratory culture and algal infection6. *Aiptasia* larvae, in particular, are critical to the study of coral symbiosis because the larval stage is the earliest development stage in which cnidarians can take up algal symbionts(2, 6, 10).

To date, live imaging of *Aiptasia* larvae has proven experimentally intractable because the larvae are highly motile, complicating observation of symbiosis induction, stasis, and bleaching within a single larva over time11. Previous studies have relied on adult or larval fixation, which reveals static algal presence and distribution, but prevents dynamic observation of the host-symbiont relationship5. We therefore sought to develop an easy-to-use device to enable live observation of *Aiptasia* and their algal symbionts over long time courses (2-12 hours) and under different environmental stressors.

Microfluidic devices have served as excellent platforms for similar long-term imaging tasks in mammalian cells(12, 13), bacteria(14–16), and plants(17, 18). Indeed, a ‘coral-on-a-chip’ microfluidic platform was previously developed for bulk reef-building coral microscopy19. However, this platform imaged non-motile juvenile coral, which passively settled in the device, and cannot be translated to the study of highly motile larvae. To our knowledge, no published study has demonstrated live imaging of *Aiptasia* larvae (or other similar cnidarian larvae) and their associated algal symbionts, likely due to the challenges associated with designing structures to immobilize deformable, highly motile cnidarian larvae that are ~100 μm in size.

Here, we present a single-layer microfluidic device and setup scheme for isolating, trapping, and imaging up to 90 live, individual *Aiptasia* larvae and their algal symbionts in parallel. We designed the device as part of a university microfluidics laboratory course that brought together engineers and life scientists to find solutions for pressing biological problems. The device is designed for operation by life scientists unfamiliar with microfluidics, requires only simple and inexpensive equipment, and is compatible with standard cnidarian-rearing protocols.

Optimizing the device required careful consideration of multiple trap geometries and organism sizes. To test how trap and organism geometries affect trap array performance, we quantified trap occupancies for polymer beads of various sizes in devices of different heights and trap apertures. To ensure that adequate nutrients would reach trapped organisms, we simulated flow fields within and around occupied and unoccupied traps. Using an optimized trap design for *Aiptasia*, we performed live imaging with *Aiptasia* larvae and their algal symbionts under constant nutrient flow and demonstrated the device’s potential for bleaching studies via the capture of an algal expulsion event under the environmental stressor DCMU (3-(3,4-dichlorophenyl)-1,1-dimethylurea, known by its trade name Diuron). These experiments highlight the utility of the device for studying early cnidarian symbiosis and investigating the mechanisms of stress-driven coral bleaching. Moreover, our modeling and bead-loading results provide critical information for future adaptation of these devices for study of a wide range of large, deformable, and motile organisms.

## Results and Discussion

### Microfluidic Device Design and Experimental Setup

A microfluidic device capable of studying anemone larval-algal symbiosis and mechanisms of algal expulsion must satisfy several design criteria. The device must stably trap deformable and motile *Aiptasia* larvae at medium- to high-throughput (10-50 larvae per experiment) over multiple hours (2-24 hours) while retaining organism viability. In addition, the device must allow for sufficiently high spatial and temporal resolution in both transmitted light and fluorescence channels (for imaging algal autofluorescence) to image larvae (~90-140 μm), associated internal structures (*e.g.* gastrodermal layer and gastric cavity), and their algal symbionts (~5-10 μm). Finally, an ideal device for use in life science laboratories requires little setup and training time (~30 and ~15 minutes, respectively) and can be operated using common laboratory equipment (*e.g.* syringe pumps).

To meet these criteria, we designed a single-layer polydimethylsiloxane (PDMS) trapping device (termed ‘Traptasia’ for Trapping of *Aiptasia*) that requires only a syringe pump as necessary additional equipment **[Fig. 1]**. Each device contains an array of 90 paired isosceles triangle traps at 75° pitch (relative to centerline of the pair) spanning the full chamber height **[Fig. 1a-c]**; each triangle trap is 373 μm long and 200 μm wide with a variable trap aperture between triangles **[Fig. 1b]**. Traps were laterally spaced by 280 μm and arranged in 16 rows with alternating spacing to prevent clogs and promote nutrient flow between traps. Two additional traps were placed near the inlet and outlet of the device to prevent chamber collapse. The full trap array totals 20,000 μm in length and 4,000 μm in width with a single inlet and single outlet port, both designed for 23G pin to external tubing connections (*e.g.* Tygon). These devices are readily compatible with standard inverted microscopy setups present in life science laboratories **[Fig. S1]**.

**Fig. 1.**
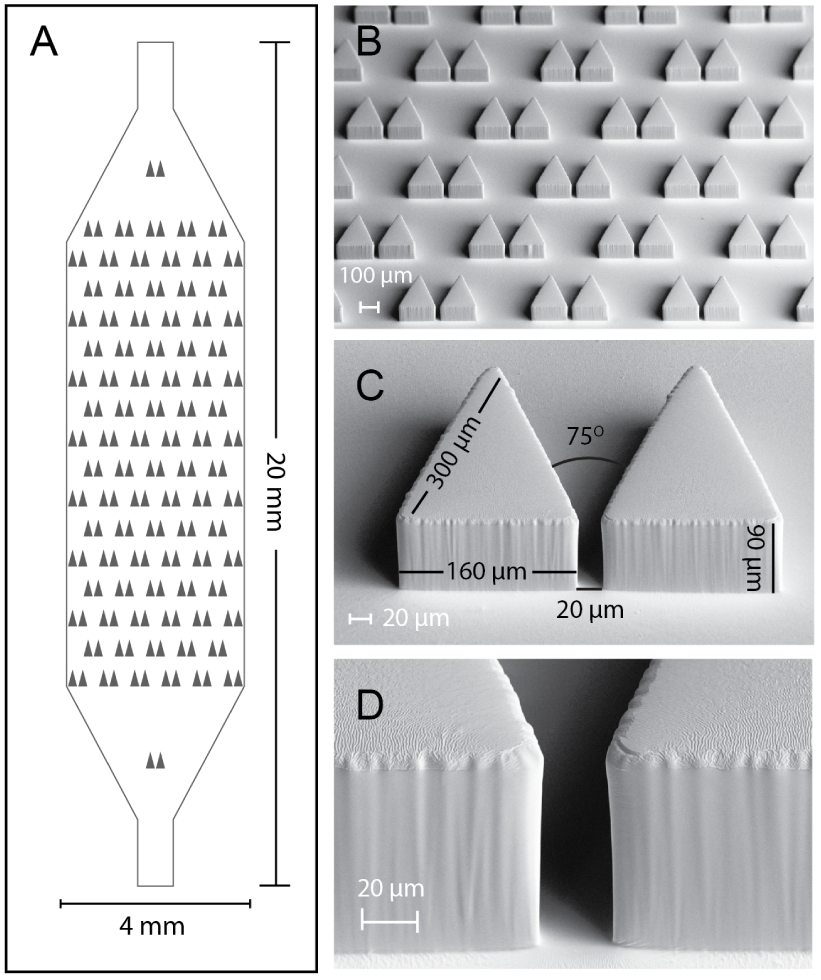
Single-layer microfluidic device for trapping individual *Aiptasia* larvae. (A) Design schematic for ‘Traptasia’ device containing an array of 90 triangular traps and tubing inlets. The final PDMS device is sealed to a No. 1 coverslip for imaging. (B) SEM image of PDMS device showing trap array. (C) SEM image of a single trap with labeled dimensions. (D) SEM image of trap aperture walls.

### Validation of the Trapping Array for Large Cell and Organism Loading

To enable medium-to high-throughput imaging of trapped organisms, specimens must be efficiently loaded in the device trapping array. To assess trap loading efficiencies, we systematically measured trap occupancy for different trap geometries using polymer beads as a proxy for organisms of different sizes. Polystyrene beads (~5000 beads per condition, similar to projected specimen concentrations) of ~ 40, 80, and 100 μm mean diameters were loaded into ‘Traptasia’ devices with trap apertures of 20, 30, 40, and 50 μm and chamber heights of 50, 70, and 90 μm. Trap occupancy across the trapping array was determined after flow stabilization at 100 μL/min **[Fig. 2a]**. Beads larger than the chamber height were unable to reliably enter the device (as confirmed by imaging beads at the device inlet) and were excluded from further analysis (grey asterisks, **[Fig. 2a**]).

**Fig. 2.**
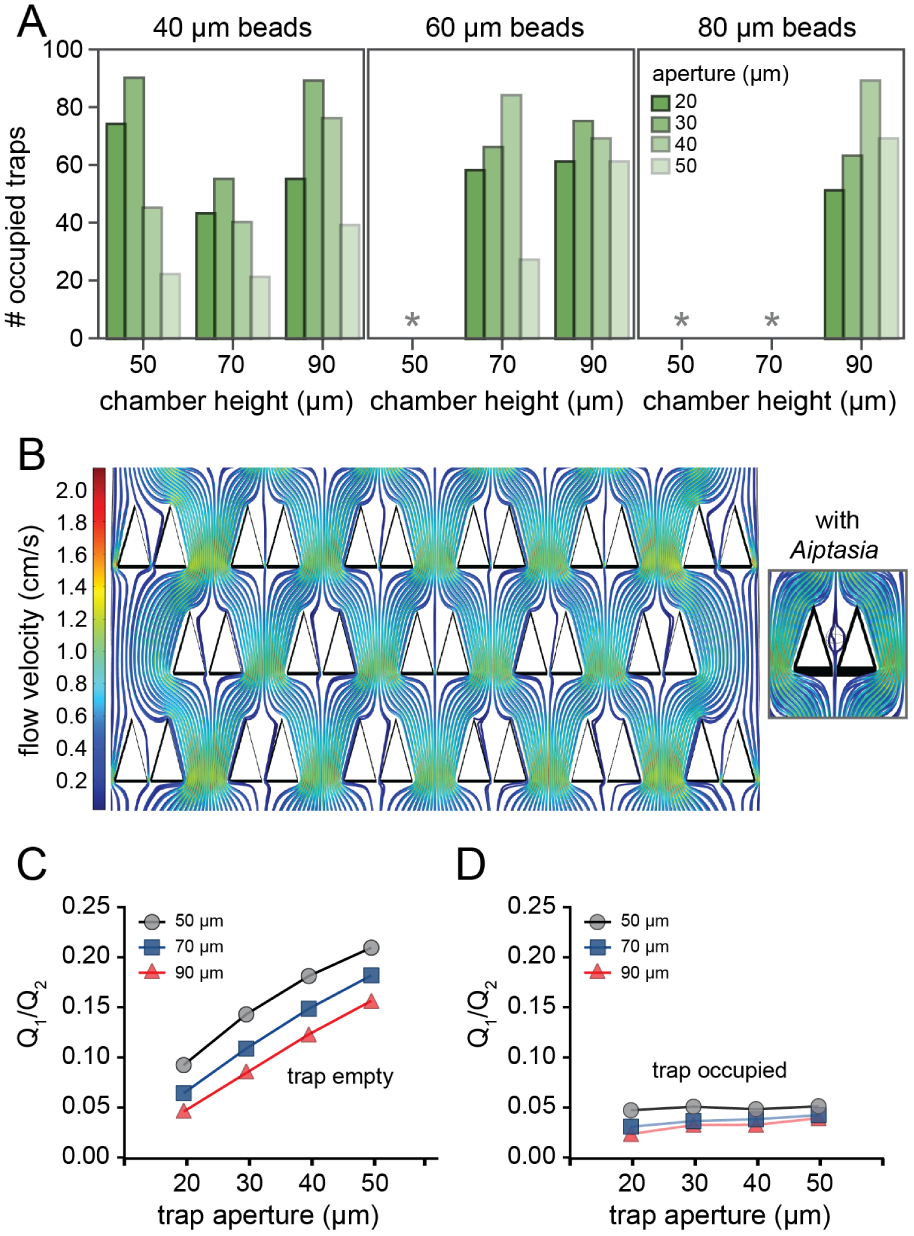
Trap-loading efficiencies and fluid flow simulations. (A) Bead occupancy histograms from bead loading experiments with different trap apertures, chamber heights and bead sizes. Grey asterisks denote instances when beads were too large to consistently enter chambers. (B) Simulated flow profiles within and between unoccupied traps under laminar flow conditions. Inset shows flow profiles around a rigid body as a proxy for an *Aiptasia* larva. (C,D) Ratio of simulated flow within and between traps (Q1/Q2) when unoccupied (C) and (D) occupied by simulated *Aiptasia* larvae for 3 chamber heights.

*Aiptasia* larvae are highly deformable, suggesting that geometries that efficiently trap beads with a wide size range (*e.g.*, 40 - 100 μm) would work best. Small beads (~40 μm) were trapped efficiently at all chamber heights at narrow trap apertures (20, 30, 40 μm), but often escaped through, or around, traps at the largest aperture traps (50 μm). This trend persisted for 60 and 80 μm beads in taller devices (70, 90 μm chambers), but narrow trap apertures (20, 30 μm) yielded slightly lower occupancies. This likely results from increased fluidic resistance at lower trap apertures, as well as an observed higher propensity for clogging at device inlets. From these data, we estimated that *Aiptasia* larvae of size range ~40-100 μm under deformation should achieve trap occupancies of 20 - 80 traps loaded (of 90 potential traps) per experiment when ~5,000 larvae (approximate medium spawn size) are loaded. These experiments represent an estimate of feasible organism loading; smaller spawn inputs may result in less efficient trap loading but should follow similar relative distributions to those reported here. Furthering this claim, we noted during loading experiments most traps were occupied within the first ‘burst’ of bead flow (~10 s, <1000 beads), suggesting that excess loading concentrations may not linearly increase occupancy.

Based on these results, we tested loading of *Aiptasia* larvae into trap arrays with 90 μm tall chambers and 20 μm trap apertures and found this geometry excellent for anemone trapping. Differential bead counts at this geometry also suggested single-organism-loading would be feasible **[Fig. S2]**. Unlike polymer beads, *Aiptasia* larvae are highly deformable and motile (motility in culture shown in **[Movie S1]**) and therefore require constant fluid flow to keep them stably trapped. Flow rates of 35-100 μL/min largely held larvae in traps without causing them to squeeze through narrow trap apertures; we subsequently modeled nutrient fluid flow to investigate if larvae would remain viable during high flow rate trapping.

### Simulations of Nutrient Delivery to Trapped Aiptasia

Long-term (2-24 hour) imaging of *Aiptasia* larvae requires that larvae receive adequate nutrient flow to maintain viability. Within the laminar flow regime in microfluidic devices, passive diffusion is minimal; therefore, we reasoned there must be adequate flow within traps to deliver nutrients. To probe effects of trap occupancy on nutrient delivery, we simulated fluid flow through unoccupied or *Aiptasia*-occupied traps (modeled as a rigid body) and calculated the relative fluid flow ratio (Q1/Q2) between (Q1) and through (Q2) traps at 100 μL/min using COMSOL **[Fig. 2b-d]**. All geometries had non-zero Q1/Q2 values, indicating sufficient nutrient flow through traps. In particular, the 90 μm chamber height with a 20 μm trap aperture resulted in a Q1/Q2 = 0.046 for an unoccupied trap and Q1/Q2 = 0.025 for rigid ellipsoid occupied trap. The latter creates a lower bound for nutrient flow in the case of a highly non-deformable cell type; *Aiptasia* likely experience increased nutrient flow due to flow-induced deformation.

We next simulated flow profiles throughout a device with 50% occupied traps (distributed randomly). Flow through these traps (occupied or vacant) at a volumetric flow of 100 μL/min ranged between 0.0132 and 0.005 μL/s. Even at this lower bound, traps would receive 1 *Aiptasia*-volume worth of flow (0.0006 μL) in less than 1 second, sufficient for nutrient delivery and long-term incubation in the device. These simulations also revealed that flow rates were relatively uniform for different traps in the array even under randomly distributed occupancies **[Fig. S3]**. Taken together, these data establish that fresh seawater should reach and sustain trapped organisms.

### ‘Traptasia’ Enables Live Imaging of *Aiptasia* Larvae and Algal Symbionts for Long Time Courses

To perform long time-course imaging of trapped larvae, we mounted the ‘Traptasia’ microfluidic device on a spinning disk confocal microscope. We loaded larvae co-infected with algae into the device and imaged each occupied trap in the array at 40 minute intervals and at multiple heights (3 μm stepped z-stacks) under a constant seawater bath to monitor the larval-algal symbiotic relationship.

Images of larvae via brightfield microscopy clearly resolved animal margins and internal larval anatomy including the gastroderm, gastric cavity boundaries, cilia, and the apical tuft **[Fig. S4]**. Simultaneously acquired fluorescence images in a custom far-red fluorescence channel corresponding to chlorophyll autofluorescence allowed visualization of algal symbionts within the *Aiptasia* larva gastroderm (false colored to blue-green, **[Fig. S4]**).

In multiple experiments, we obtained images of >100 individual *Aiptasia* larvae, demonstrating stable trapping within the ‘Traptasia’ array. In one experiment, 25 of 90 traps remained occupied by larvae (each with 5-30 algal symbionts) under 35 μL/min seawater flow for >10 hours (example traps, **[Fig. 3]**). Of these larvae, all 25 remained viable for the duration of the 10 hour experiment (**[Figs. S5, S6]**). After flow was terminated at ~11.5 hours, 8 larvae swam out of the trap (TrapIDs: 23-32, 35), and 1 underwent cell lysis (TrapID: 26) **[Fig. S7]**, demonstrating that high flow rates are necessary for stable trapping.

**Fig. 3.**
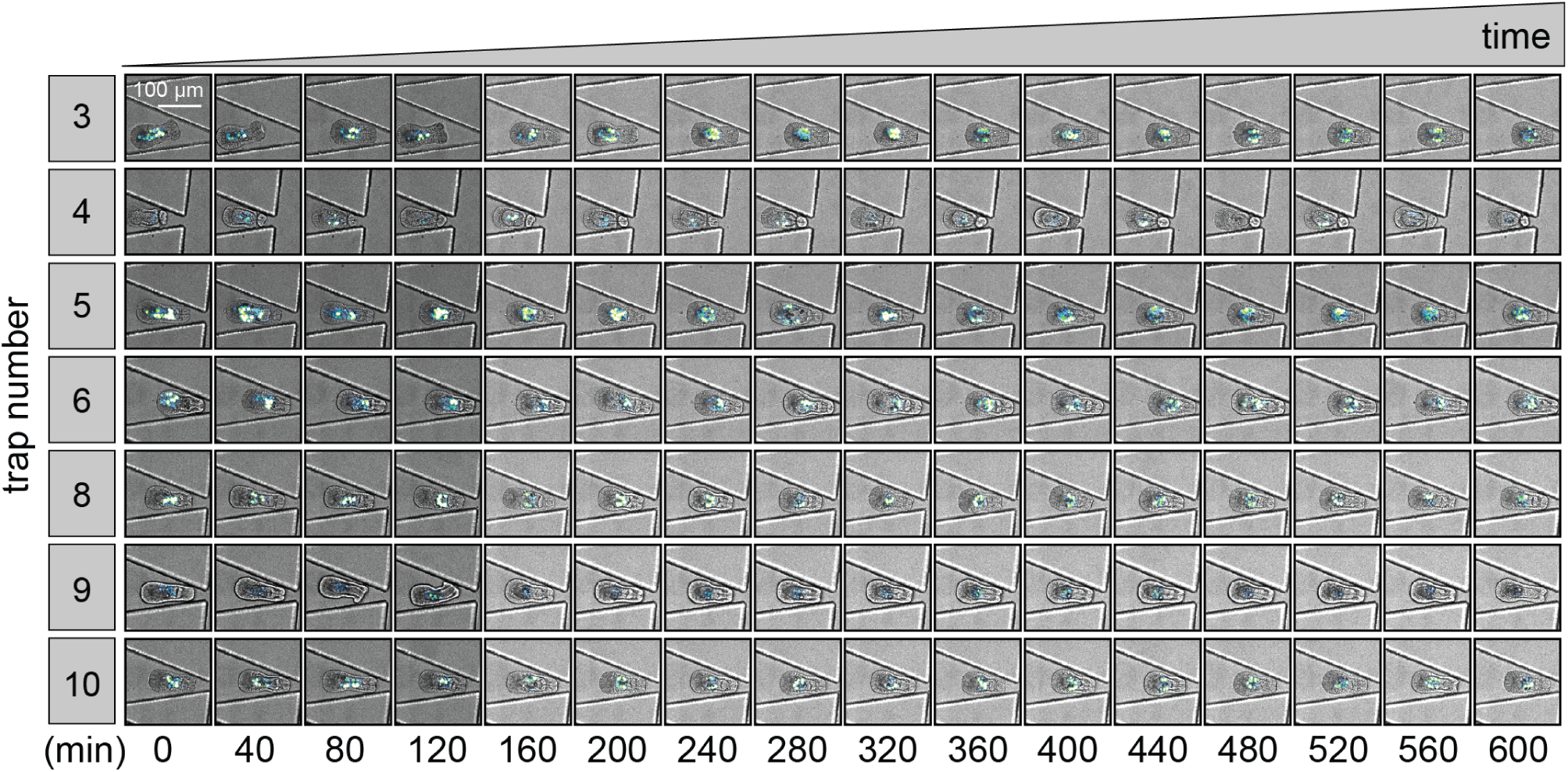
Time course of select traps demonstrating stable trapping of viable *Aiptasia* larvae for 10 hours. Each row represents a single trap (TrapID, left); each column represents an imaged time point. Each image is a merged brightfield and fluorescence image to show device trap, trapped larva, and algal symbionts. All traps shown in **[Figs. S5, S6]**.

### Detailed Algal Symbiont Imaging in Individual Larvae

The ability to visualize and track individual algal symbionts within live larvae during homeostasis and under stress is essential to understand the mechanisms of cnidarian-algal symbiosis, as well as aberrant algal expulsion associated with reef bleaching. However, current algal tracking methods require extensive confocal or electron microscopy on fixed samples and thus cannot provide dynamic information about algal movements5. Trapped *Aiptasia* larvae were viable with free-moving cilia in ‘Traptasia’ devices with 90 μm chamber heights but appeared slightly compressed, suggesting that the devices could stabilize motile larvae for algal symbiont imaging **[Movies S2]**.

To track individual algae, we trapped co-infected larvae and algal symbionts and performed fast-acquisition imaging in transmitted light alone (50 ms exposures, 5 ×3 μm z-stacks) to obtain continuous time series for select, individual *Aiptasia* larvae. Under normal seawater conditions at 100 μL/min flow, algae appeared to freely move within the gastroderm, driven mostly by organism spin, without entering the gastric cavity**[Fig.4]**. Algae counts per organism remained constant throughout acquisition. Larval revolutions, including change in direction and speed, were clearly visible, providing additional opportunities to monitor complex or subtle stress responses **[Movies S3, S4]**. The ability to resolve individual algae within larvae without confocal microscopy could allow use of the device within a broad range of environments (*e.g.* undergraduate research institutions, local museums or aquaria, or field research stations).

**Fig. 4.**
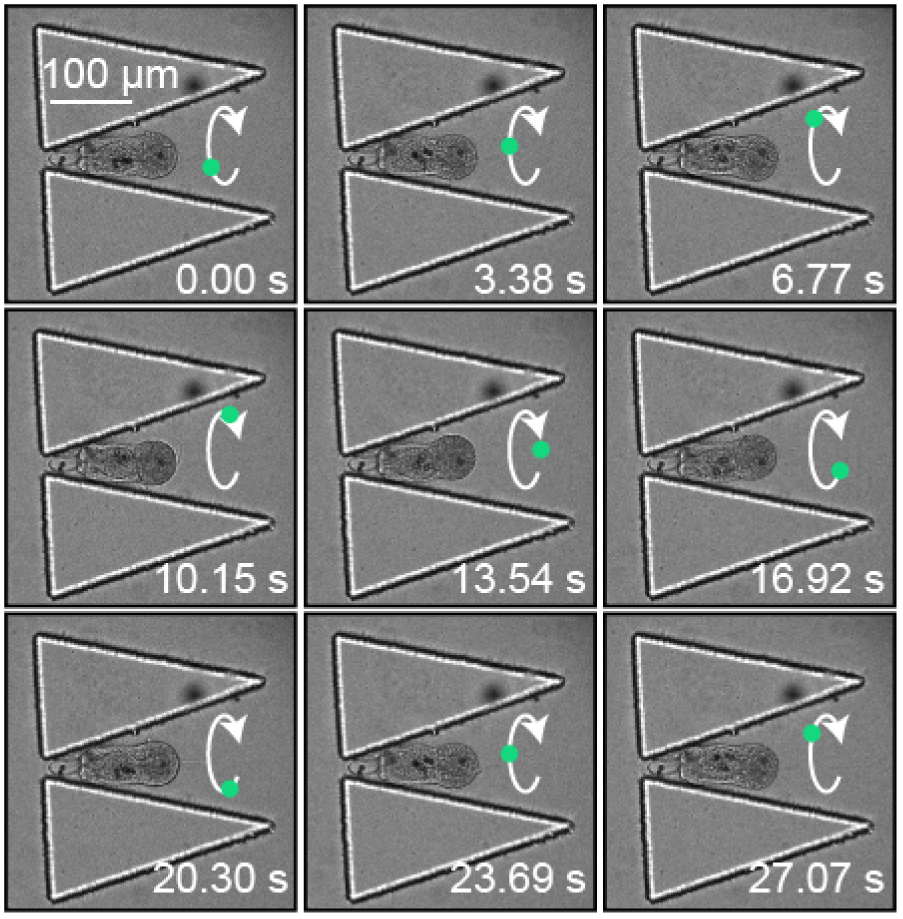
Brightfield imaging of a trapped *Aiptasia* larva at high temporal resolution demonstrating the ability to resolve algal movement and monitor larval revolutions (larval spin, relative to lengthwise normal, denoted by white arrow; estimated position in revolution denoted by green dot) within the trap. Frames were extracted at a single plane every 5 time steps (for visualization) from a continuous time-lapse sequence.

### Multiple Larval-death Mechanisms Revealed by Imaging

Viable *Aiptasia* larvae show robust ciliary movement and low-speed rotation. During our trials, we observed multiple events of larval death characterized by high rotation within traps, lack of ciliary movement, or cellular degradation **[Fig. S8]**. Bubble nucleation within the microfluidic device during initial trials often resulted in larval death **[Fig. S8c]**, but was mitigated entirely with at 100 μL/min flow speeds in combination with device debubbling (see *Supplemental Information*).

Beyond device-related larval death phenotypes, we observed interesting, potentially physiologically-relevant instances of organism degradation. In some cases, dying larvae separated their mouth from their gastric cavity prior to complete degradation **[Movie. S5]**, a phenomenon that has also been observed in our larval cultures. In other cnidarian species, mouth separation may precede opening for feeding or organism regeneration20, but these mouth separation events merit further study in *Aiptasia* to determine physiological significance. In other cases, larval degradation appeared to result from ciliate infestation, likely acquired during algal infection **[Fig. S8b]**. These events are frequently observed in bulk culture but have never been visualized in cellular detail, illustrating how the ‘Traptasia’ device can facilitate detailed investigation of multiple aspects of cnidarian physiology.

### Observation of Algal Symbiont Explusion from a Larval Host under a Proposed Environment Stressor

To investigate the cellular mechanisms that drive cnidarian bleaching, we introduced a proposed coral stressor, DCMU (3-(3,4-dichlorophyli)-1,1-dimethylurea, trade name Diuron) to trapped *Aiptasia* larvae and their algal symbionts. In the presence of 25 μM DCMU in seawater at 35 μL/min flow rates, 33 *Aiptasia* larvae and their algal symbionts were stably trapped within 90 possible trapping positions **[Fig. S9]**. Of the 33 trapped larvae, 28 remained trapped for the course of the imaging acquisition (80 minutes). Five larvae ‘swam’ upstream of flow or through the trap aperture (Trap IDs: 8, 10, 11, 29, 42), suggesting that larval motility and deformation may increase under stress as compared to seawater.

Under DCMU treatment, we captured an algal expulsion event from a live *Aiptasia* larva **[Fig. 5a, Movie S6]**. Quantification of mean algal autofluorescence within larval margins and the surrounding external environment from sum projection before, during, and after the expulsion of algal symbionts from the larval gastric cavity and mouth reveals a strong signal gain during expulsion in the external environment and consequent loss within the host margins. This analysis provides a high-throughput method for identifying expulsion events without manual inspection **[Fig. 5b]**. To our knowledge, these data represent the first dynamic visualization of an algal expulsion event, thought to be the critical event underlying widespread coral bleaching. In future implementations of the device, systematic trials of larval imaging under different stressors could capture more expulsion events and further elucidate biological mechanisms behind larval stress induction and algal expulsion.

**Fig. 5.**
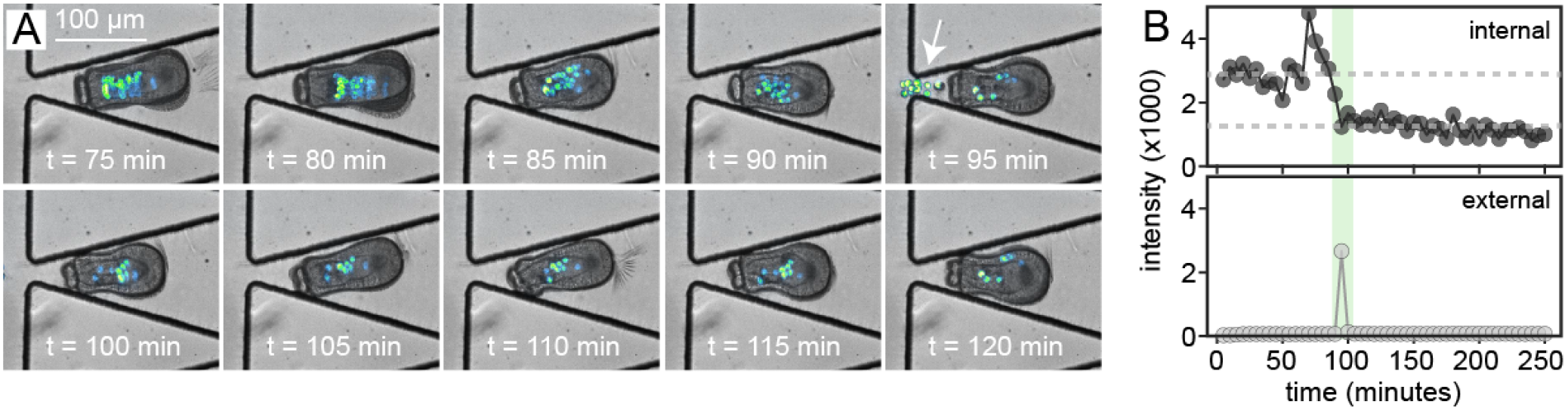
Algal expulsion event from a live larva. (*A*) Time-lapse imaging sequence showing clear evidence of an algal expulsion event from the larval mouth through the gastric cavity (white arrow, 95 minutes). (*B*) Mean measured fluorescent intensity (photon counts) for a region of interest within (top panel) and external to (bottom panel) *Aiptasia* boundaries. Fluorescence within *Aiptasia* decreases by about one third at the same frame that expelled algae are visible outside the larval body.

## Conclusions

In this report, we developed an easy-to-operate microfluidic device and setup (’Traptasia’) for trapping and imaging *Aiptasia* sea anemone larvae and their algal symbionts over long time courses. We demonstrate the utility of the device both in high-throughput imaging and detailed time-lapse imaging tasks. This device addresses a long-standing limitation in the ability to image live, highly motile larvae for mechanistic studies of the processes that drive coral reef bleaching.

In confirming the viability of the ‘Traptasia’ device design, we conducted bead occupancy tests across a wide array of trap geometries from trap apertures of [20, 30, 40, 50] μm and chamber heights of [50, 75, 90] μm. These occupancy experiments were used identify device geometries for successful *Aiptasia* trapping, but they could easily be used to estimate optimal trap parameters and likely trapping occupancies for other large and highly motile organisms. While prior data exist for mammalian single cell traps, optimal trapping geometries and predicted occupancies for large (>30 μm size) organisms are not readily available.

Similarly, our results presented in flow field calculations across different device geometries also have utility outside device parameterization for these studies alone. In future work, these flow profiles will help inform environmental stressor concentration dosing and resultant effective agent concentrations experienced by each *Aiptasia* larva. Additionally, this flow modeling approach may help inform nutrient concentration requirements for organisms requiring more complex nutrient solution delivery to sustain long-term study.

In the future, this device could facilitate the study of *Aiptasia*-algal symbiosis both under healthy conditions and in the presence of known and proposed inducers of coral bleaching (*e.g.* increased temperature, increased salinity, DCMU and other chemicals). The device could also help answer basic mechanistic biological questions about the stability of algal symbiosis, the frequency of healthy vs. non-healthy expulsion events, and organism-to-organism heterogeneity in symbiotic behavior (i.e., migration and algal movement). Combined with reverse-flow techniques for retrieving the larvae post-analysis, additional downstream assays such as genomic8 or transcriptomic(7, 9) profiling could help identify algal stress mechanisms, if imaging data from this platform reveals interesting larval behaviors to a given stressor.

This device continues to be used for mechanistic biological studies of cnidarian-dinoflagellate symbiosis in collaborating laboratories at Stanford and the Carnegie Institute for Science. In future work, we hope the device can be adopted more broadly, both for experiments involving sea anemones, corals and other cnidarians, as well as other large, motile organisms. By sharing these data, design files, and detailed protocols, we hope that other biologists and researchers will adopt this, or similar, trapping devices for their own organism research.

## Materials and Methods

### Animal Culture and Maintenance

Small sea anemone (*Aiptasia*) larvae were cultured in the laboratory21. Larvae were obtained via induced spawning of clonal adult *Aiptasia* anemone lines CC7 (male) and PLF3 (female) in paired co-culture in artificial sea water (ASW; 34 ppt Coral Pro Salt in deionized (DI) water, pH 8.0, Red Sea Inc.)9. To set up anemones for spawning, adults from each clonal population were selected by size (>15 mm disk size) and manually paired (single male to single female) in polycarbonate tanks of 300 mL of ASW to induce spawning under 27°C incubation (Percival). Co-cultures were maintained on a 12-hour light: 12-hour dark cycle with 25 μmol photons/(m^2^ s) white fluorescent light (Philips ALTO II 25W) (light cycle) for 25 days and subsequently switched to a 16-hour light: 8-hour dark cycle for 5 days, with an additional 1 μmol photons/(m^2^ s) actintic blue LED (Current USA TrueLumen) during the dark cycle. One-day-old larvae were re-suspended in 1 mL ASW in wells of a 24-well polypropylene (PPE) plate at a density of 1,000 - 3,000 larvae per well before infection with algae.

### Algal Infection for Symbiosis Studies

Freshly isolated algae were used to infect *Aiptasia* larvae in this study. *Symbiodinium* clade A were isolated from an individual CC7 adult anemone by homogenization (PowerGen 125 rotor, Fisher Scientific) at 30,000 rpm for 10 s, restrictive mixing via a 25G needle attached to a 1 mL syringe (BD Biosciences) at 5 repetitions and centrifugation at 1000 ×g for 2 min to pellet algal cells. Intact algal cell pellets were washed 10 times in fresh ASW and re-suspended 1:1 v/v in wells of the 24-well PPE plate containing larvae. The plate was maintained on a 12-hour light: 12-hour dark cycle at 27*°*C prior to use in imaging studies in the microfluidic device platform within 1-2 days of infection. Larvae were 3-5 days old at the start of the experiment.

### Device Design, Photolithography, and Fabrication

Devices of varying trap aperture and chamber height were fabricated via standard soft lithography protocols22. Design files used in this study are provided as **Supplemental Resource 1**. Full lithography details are provided in **[Table S3]** and **[Fig. S10]**.

### Bead Loading Experiments

Different sizes of polystyrene cross-linked polymer beads (40, 60, 80 μm general distribution, products PPX-400, PPX-600, PPX-800, Spherotech) were assessed in the ‘Traptasia’ device. Using a similar setup to that used for *Aiptasia* experiments, per each condition ~5000 beads at constant volume were loaded for ~10 s, the device was subsequently de-bubbled, and loading was continued and allowed for stabilize for 2 minutes under 100 μL/min flow (infuse-only) using a syringe pump (Pico Pump, Harvard Apparatus).

### COMSOL Fluid Modeling

Fluid flow simulations to assess flow fields and model nutrient flow through traps with and without *Aiptasia* were modeled in COMSOL (COMSOL Multiphysics) using the 3D Laminar Flow module. Inlet velocity was set at 100 μL/min and outlet pressure was set to 1 atm. *Aiptasia*-occupied traps were simulated by placing a rigid ellipsoid of volume 5.9 *10^5^ μm^3^ 1 μm away from the trap aperture center for trapped flow field analysis. Further parameters are described in **[Table S-T1]**.

### Device Loading of *Aiptasia* Larvae

To perform *Aiptasia* imaging experiments, *Aiptasia* larvae were loaded directly from culture wells (~100 uL) using 1/16" tubing (Tygon, Fisher) attached to a 1 mL syringe (Plastipak, BD) with a 23G luer-lock connector (McMaster-Carr). Lines were hung from the syringe pump vertically for 2 minutes to concentrate the motile larvae and connected upstream of the inlet stopcock. A loading protocol is provided in *Supplemental Information*. In trials with DCMU treatment, seawater containing 25 mM DCMU (Diuron, Sigma) was introduced instead of normal seawater after larval loading.

### Image Acquisition and Analysis

Four-dimensional imaging data (time, fluorescence channels, and z-slices) of *Aiptasia* larvae in the microfluidic device traps were acquired on a Leica DMI6000B stand equipped with a Yokogawa CSU-10 spinning disk head, QuantEM camera (Photometrics), and ASI MS2000 motorized XY stage under 20X magnification (Leica 20X/0.7 NA multi-immersion, used with glycerine solution). Z-stacks were collected in transmitted light and chlorophyll autofluorescence channels (561 nm excitation, 405/488/561 nm dichroic and 637/37 nm emission, Semrock), as noted, under control of SlideBook 6 (Intelligent Imaging Innovations). Images were analyzed in Fiji (ImageJ)23.

### Open Data Access and Supplemental Information

An open-source project repository (containing design files, imaging data for all trials (compressed as .avi image sequences), protocols, and additional resources, as mentioned in the text) is available at our **Open Science Framework** page, which can be found at OSF under [doi:10.17605/osf.io/j2rsy]. Extended methods, protocols, and troubleshooting notes are available in *Supplemental Information*.

## ACKNOWLEDGMENTS

This project originated during a 10-week Microfluidic Device Laboratory course at Stanford (BIOE 301D/GENE 207, W17) and we acknowledge departmental support (Stanford Department of Bioengineering, Genetics) for course resources and fabrication time. Devices were fabricated in the Stanford Microfluidics Foundry. *Aiptasia* were reared and kindly provided by the laboratory of J. Pringle, with assistance of A.Formica and O. Barry. P.M.F. is a Chan Zuckerberg Biohub Investigator and wishes to acknowledge the support of the Alfred P. Sloan Foundation. K.K.B. acknowledges support of a Chem-H Chemical Biology Interface (CBI) Training Grant (Stanford University). W.V.T and K.K.B acknowledge the support of NSF GRFP Graduate Fellowships.

